# Distinct associations between thalamo-amygdala pathways and state anxiety

**DOI:** 10.64898/2026.06.08.730327

**Authors:** Martina T. Cinca-Tomás, Emmanouela Kosteletou-Kassotaki, Judith Domínguez-Borràs

**Affiliations:** Brainlab-Cognitive Neuroscience Research Group, Department of Clinical Psychology and Psychobiology, University of Barcelona, Barcelona, Spain (08035); Institute of Neurosciences, University of Barcelona, Barcelona, Spain (08035)

**Keywords:** thalamus, pulvinar, medial geniculate body, amygdala, state anxiety, subcortical low roads, diffusion MRI

## Abstract

Neurobiological models of emotion have proposed the existence of multiple direct subcortical pathways in humans, often referred to as “low roads”, linking the thalamus to the amygdala and implicated in affective function. Among these, pulvinar-amygdala structural connectivity has been associated with individual differences in anxiety and anxiety-related conditions. However, whether distinct thalamo-amygdala pathways across thalamic subnuclei differentially relate to anxiety remains unknown. Using diffusion MRI in 34 healthy participants, we reconstructed four candidate subcortical “low roads” bilaterally from the medial geniculate body (MGB), as well as the medial, inferior and lateral pulvinar to the basolateral amygdala (BLA). We then tested whether their structural connectivity strength was associated with individual differences in state and trait anxiety. Linear regression analyses revealed that fiber density in three left thalamo-amygdala pathways predicted state, but not trait, anxiety. Importantly, our results showed a functional dissociation across pathways. While fiber density in MGB-BLA and medial pulvinar-BLA pathways was negatively related to state anxiety, the inferior pulvinar-BLA tract showed the opposite association. These findings support differentiated contributions across thalamo-amygdala pathways in humans to state anxiety. The results highlight these subcortical pathways as potentially relevant neurobiological substrates for understanding anxiety and affective function.

**Key points:**

- Fiber density of three left thalamo-amygdala pathways explained 24.1% of the variance in state anxiety across 34 healthy individuals
- Fiber density in the left medial geniculate body and left medial pulvinar-amygdala pathways was negatively associated with state anxiety
- Fiber density in the left inferior pulvinar-amygdala pathway was positively associated with state anxiety

## Introduction

The ability to process affective information efficiently is a critical evolutionary adaptation across vertebrates. In humans, this ability remains essential for adaptive behavior, facilitating rapid affective responses to potential threats. Extensive evidence across non-human species has identified several subcortical neural pathways, or “low roads”, supporting affective processing under different experimental conditions through direct anatomical connections between the sensory thalamus and the amygdala [Carr, 2015; LeDoux et al., 1984; LeDoux, 2000; McFadyen et al., 2020; Phelps and LeDoux, 2005]. In particular, the basolateral amygdala (BLA), a key structure in fear conditioning and other emotion-related processes, has been identified as a primary target of thalamic projections involved in affective function [Janak and Tye, 2015; Phelps and LeDoux, 2005].

In humans, research on the subcortical amygdala pathways has predominantly focused on superior colliculus (SC)-pulvinar-amygdala connectivity, particularly in the context of visual affective processing [Carr, 2015; Kragel et al., 2021; McFadyen, 2019; McFadyen et al., 2019; McFadyen et al., 2020; Pessoa and Adolphs, 2010; Tamietto and De Gelder, 2010]. This pathway has been proposed to support rapid transmission of visual information to the amygdala, and has been implicated in several fear-related processes, including non-conscious fear processing in blind patients with bilateral occipital damage [Tamietto and De Gelder, 2010] and multiple fear-related behaviors (see [Celeghin et al., 2015; Domínguez-Borràs and Vuilleumier, 2022; McFadyen et al., 2020; Vuilleumier, 2005]). In the healthy population, white matter microstructure along this pathway predicts individual differences in attentional biases towards threatening visual stimuli [Koller et al., 2019]. Notably, structural and functional alterations within this specific pathway have been associated with different anxiety-related conditions, including phobia, post-traumatic stress disorder (PTSD) and social anxiety [McFadyen et al., 2020]. For instance, pulvinar-amygdala connectivity strength is associated with symptom severity in specific phobias [Nakataki et al., 2017]. Similarly, altered functional connectivity between low-level emotion-related regions (e.g. SC) and amygdala, as well as BLA reactivity to non-conscious fearful stimuli (typically associated with pulvinar-amygdala connectivity) have been observed in PTSD and high-anxiety cohorts [Bishop et al., 2004; Etkin et al., 2004; Neumeister et al., 2018; Steuwe et al., 2015], suggesting that altered subcortical amygdala connectivity may contribute to the anxiety-related symptomatology. However, pulvinar-amygdala connectivity has primarily been investigated considering the pulvinar as a broad thalamic structure, despite evidence that it comprises multiple anatomically and functionally differentiated subdivisions [Bourgeois et al., 2020; Froesel et al., 2021; Froesel et al., 2024]. While some pulvinar nuclei have been predominantly associated with visual and visuospatial processing, others may additionally contribute to multisensory and auditory-related functions [Froesel et al., 2021; Froesel et al., 2024]. In recent work with diffusion magnetic resonance imaging (MRI) tractography, we identified several robust and anatomically differentiated pathways linking the medial, inferior and lateral pulvinar nuclei with the BLA in humans, with potentially independent trajectories and connectivity profiles [Kosteletou-Kassotaki et al., 2026b].

In parallel, additional subcortical pathways may also play a critical role in affective processing. In particular, we have recently reported strong connections between the medial geniculate body (MGB), the main auditory thalamic relay, and the BLA, with structural connectivity of this pathway showing associations with individual fearfulness [Kosteletou-Kassotaki et al., 2026a] and amygdala responses to fear-associated sounds [Cinca-Tomás et al., 2025]. These findings are consistent with extensive animal evidence supporting the role of MGB-BLA circuitry in auditory affective processing [Keifer et al., 2015; LeDoux et al., 1984; Romanski and LeDoux, 1992], mediating fear responses across species [Khalil et al., 2023; Shinonaga et al., 1994]. However, the association of this pathway with anxiety levels in humans remains undetermined (see Rafal and Koller, 2025).

Overall, several subcortical thalamo-amygdala pathways have been identified in humans, with potentially diverse contributions to affective function. However, whether these subcortical pathways differentially relate to anxiety variability remains unknown. Here, we investigated whether structural integrity within these subcortical amygdala pathways relates to individual differences in anxiety in healthy participants. Specifically, we reconstructed bilateral thalamo-amygdala tracts between the MGB and the BLA, as well as between the medial, inferior and lateral pulvinar nuclei and the BLA, and then tested whether fiber density of these tracts predicted individual state and trait anxiety measures.

## Materials & Methods Subjects

Thirty-seven healthy participants, mostly recruited among University of Barcelona students, provided informed consent to participate. All were right-handed, reported normal hearing and no history of neurological or psychiatric disorders. One participant was excluded from the analysis due to low signal-to-noise ratio in the MRI data and drowsiness during task performance, and two additional participants were excluded following visual inspection, as they presented extreme values in tract-specific fiber density measures. Thus, the final sample consisted of thirty-four volunteers between 18 and 39 years of age (18 females; mean age = 24.74). All procedures were approved by the ethics committee of the University of Barcelona (RB00003099 - CER042405), in accordance with the Declaration of Helsinki. All participants received monetary compensation for their participation.

### Procedure

#### State-Trait Anxiety Inventory - STAI

Before starting the MRI acquisition, participants completed the validated Spanish version of the State-Trait Anxiety Inventory (STAI; [Spielberger et al., 1982], a 40-item self-report questionnaire divided in two parts assessing state (STAI-S) and trait anxiety (STAI-T) respectively. For the STAI-S (20 items), participants were instructed to rate their current feelings of anxiety on a scale from 0 (not at all) to 3 (very much so). For the STAI-T (the remaining 20 items), participants indicated how frequently they generally experience anxiety-related symptoms, using the same scale from 0 (almost never) to 3 (almost always). The total score of each part may range between 0 and 60, with higher scores reflecting higher levels of anxiety (see Fig. 1). One participant completed the validated English version of the test. Since in the English version the response scale ranges from 1 to 4, scores were linearly rescaled to match the Spanish 0–3 scale prior to analysis.

**Figure 1.**
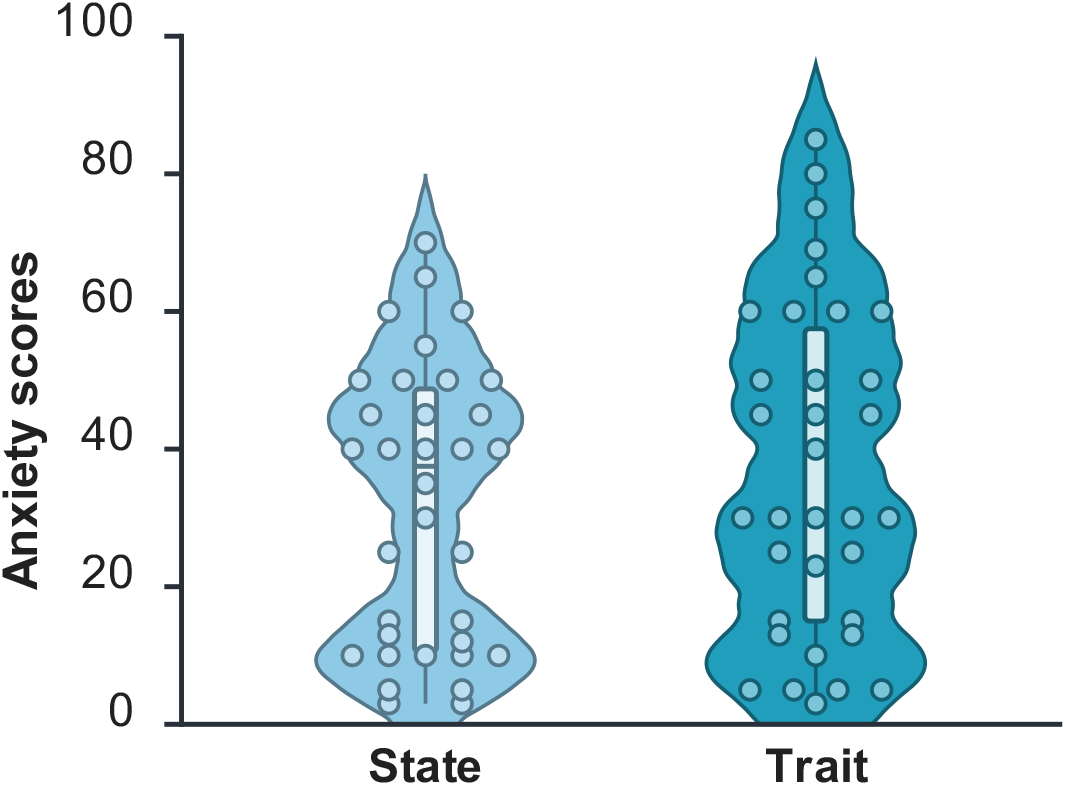
Distribution of individual state and trait anxiety scores across participants. Each dot represents an individual participant, and the central boxplot indicates the median and interquartile range. Figure created with BioRender.

### Data acquisition

Whole-brain MRI data were acquired on a 3T Philips Ingenia CX scanner equipped with a 32-channel whole-head coil, located at the BarcelonaBeta brain research center (https://www.barcelonabeta.org/es), Barcelona, Spain. The imaging protocol included T1-weighted (T1w) high-resolution structural images and diffusion-weighted imaging (DWI) sequences in a single session. The T1w structural images were obtained using a Turbo Field Echo (TFE) sequence with the following parameters: voxel size = 0.9 mm isotropic, field of view (FoV) = 240 mm, 200 slices, repetition time (TR) = 9.9 ms, echo time (TE) = 4.6 ms, and flip angle (FA) = 8°. For the diffusion-weighted imaging, a total of 108 volumes were acquired for each participant in the anterior-to-posterior phase-encoding direction. This included 96 diffusion-weighted directions distributed across multiple b-values: 8 directions at b = 500 s/mm^2^, 8 at b = 1000 s/mm^2^, 16 at b = 2000 s/mm^2^, and 64 at b = 3000 s/mm^2^. Additionally, 12 non-diffusion-weighted (b = 0 s/mm^2^) images were included in the main acquisition. For distortion and motion correction, two extra b = 0 s/mm^2^ images were acquired with opposite phase-encoding directions (anterior-to-posterior and posterior-to-anterior). DWI acquisition parameters for both the diffusion-weighted and b0 images were as follows: voxel size = 1.65 mm isotropic, 81 slices with no gap, TR = 7463 ms, TE = 89 ms, FA = 90°, and a multiband acceleration factor of 3.

### Diffusion MRI preprocessing

Preprocessing of diffusion-weighted MRI data was performed using tools from the MRtrix3 software. Denoising was first applied using the dwidenoise function, which employs a patch-based principal component analysis approach grounded in random matrix theory to leverage data redundancy [Cordero-Grande et al., 2019; Veraart et al., 2016]. Rician background noise was then removed using mrcalc. Gibbs ringing artifacts were subsequently corrected using the mrdegibbs method [Kellner et al., 2016]. Susceptibility-induced distortions and motion artifacts were corrected using FSL’s topup and eddy tools [Smith et al., 2004], as implemented within the dwifslpreproc pipeline. B1 field inhomogeneity correction was performed using dwibiascorrect, and a brain mask was generated with dwi2mask to restrict subsequent analysis to brain tissue. The T1w anatomical image was then rigidly aligned to the DWI data using ANTs.

Response functions for white matter, grey matter and cerebrospinal fluid (CSF) were estimated for each participant using the multi-shell multi-tissue constrained spherical deconvolution (MSMT-CSD) algorithm [Jeurissen et al., 2014]. These response functions were subsequently used to compute subject-specific multi-tissue fiber orientation distribution functions (fODFs), which represent possible fiber directions with corresponding weights in each voxel. Lastly, joint bias field correction and global intensity normalization of the multi-tissue compartment parameters were applied using the mtnormalise command in MRtrix3.

### Region of interest (ROI) definition

ROI definition involved processing each subject’s T1w aligned anatomical image to obtain the ROIs for tractography. This process takes the subject’s T1w image and predefined ROIs in MNI space as input, and outputs a segmented T1w image along with the ROIs in individual subject T1w space. Because the T1w image was already rigidly aligned to the DWI data, the resulting ROIs were likewise aligned and suitable for use in tractography analyses. ROI definition and anatomical segmentation procedures followed the same pipeline previously implemented in Kosteletou-Kassotaki et al. [2026b].

Cortical and subcortical segmentation of the T1w anatomical images was performed using FreeSurfer (http://surfer.nmr.mgh.harvard.edu/). The medial geniculate body (MGB), medial pulvinar (PuM), inferior pulvinar (PuInf) and lateral pulvinar (PuL) ROIs were extracted using FreeSurfer’s thalamic segmentation module, which utilizes a probabilistic atlas derived from histological data and high-resolution *ex vivo* MRI [Iglesias et al., 2018]. The basolateral amygdala (BLA) ROI was defined using the amygdala segmentation developed by Saygin et al., [2017], also implemented in FreeSurfer. The lateral, basal and accessory basal subregions were combined to form a single BLA ROI. To improve the neuroanatomical specificity of tract reconstruction, exclusion ROIs were applied based on the anatomical location and expected connectivity of the MGB-BLA, PuM-BLA, PuInf-BLA and PuL-BLA pathways (as in Kosteletou-Kassotaki et al., [2026b]). These exclusion criteria were anatomically informed following initial visual inspection and are detailed in Table 1.

**Table 1.** Tracts and ROIs included to identify them.

| Thalamus | Tract name | Inclusion ROI 1 | Inclusion ROI 2 | Exclusion ROIs |
| --- | --- | --- | --- | --- |
| **Posterior** | MGB - BLA | **MGB** | **BLA** | PuA, PuM, PuL, PuInf, MD, AV, VA, LD, VPL |
|  | PuM - BLA | **PuM** | **BLA** | PuA, PuL, PuInf, MD, AV, VA, LD, VPL |
|  | PuInf - BLA | **PuInf** | **BLA** | PuA, PuM, PuL, MD, AV, VA, LD, VPL |
|  | PuL - BLA | **PuL** | **BLA** | PuA, PuM, PuInf, MD, AV, VA, LD, VPL |

Abbreviations: MGB = medial geniculate body, BLA = basolateral amygdala, PuA = anterior pulvinar, PuM = medial pulvinar, PuL = lateral pulvinar, PuInf = inferior pulvinar, MD = mediodorsal nucleus, AV = anteroventral nucleus, VA = ventral anterior nucleus, LD = laterodorsal nucleus, VPL = ventral posterolateral nucleus

### Diffusion MRI analysis

Diffusion MRI tractography was conducted using MRtrix3, leveraging the preprocessed diffusion-weighted images together with anatomically defined ROIs to reconstruct bilateral MGB-BLA, PuM-BLA, PuInf-BLA and PuL-BLA pathways. Pathway reconstruction procedures followed a similar tractography framework to that previously employed to characterize these pathways in humans [Kosteletou-Kassotaki et al., 2026b]. Initially, whole-brain tractograms were generated using the iFOD2 probabilistic tractography algorithm, seeding 10 million streamlines with a fiber orientation distribution (FOD) amplitude cutoff of 0.06. To minimize false-positive streamlines and premature terminations, we employed the Anatomically Constrained Tractography (ACT) framework based on a five-tissue-type (5TT) segmentation of the T1w anatomical image including cortical and subcortical gray matter, white matter, CSF and pathological tissue [Smith et al., 2012]. Next, subject-specific ROI to ROI tractograms were generated by seeding 20,000 streamlines per voxel within each seeding ROI (MGB, PuM, PuInf and PuL) using the following tracking parameters: cutoff 0.12, maximum fiber length 40 mm, minimum fiber length 5 mm, FOD amplitude threshold of 0.1, and angle threshold of 30 degrees. Resulting tractograms were constrained using inclusion and exclusion ROIs (see Table 1) to ensure biologically plausible reconstructions of the respective pathways. For quantitative assessment of structural connectivity, the targeted tractograms were combined with the whole-brain tractogram, and processed using the SIFT2 algorithm. This approach assigns a weight to each streamline to achieve quantitative consistency with the underlying fiber density [Smith et al., 2015]. Streamline weights were finally multiplied by the global scaling factor μ, computed from the whole-brain tractogram, to obtain quantitatively meaningful measures of fiber bundle capacity (FBC) comparable across participants. As this measure is an index of structural fiber density [Cinca-Tomás et al., 2025; Kosteletou-Kassotaki et al., 2026b], it is hereafter referred to as fiber density.

### Relationship between fiber density of thalamo-amygdala pathways and anxiety

To explore the potential relationship between fiber density of the thalamo-amygdala pathways and anxiety, we fitted separate linear regression models for state (STAI-S) and trait (STAI-T) anxiety scores, with tract-specific fiber density as predictors [Kosteletou-Kassotaki et al., 2026a; McFadyen et al., 2019]. Concretely, eight tracts of interest were tested: MGB-BLA, PuM-BLA, PuInf-BLA and PuL-BLA from the left and right hemispheres separately. Analyses were conducted in RStudio (R Core Team, 2024). To address potential redundancy among fiber-density pathways, we assessed shared variance using principal component analysis (PCA; *princomp* function in R). To identify a parsimonious candidate regression model, we fitted a series of nested linear models with increasing numbers of thalamo-amygdala predictors. An exhaustive subset-selection procedure implemented in the *regsubsets* function from the *leaps* package was used to explore which combination of candidate predictors (tracts) best explained variability in anxiety scores. For each model size, the subset minimizing residual sum of squares was identified. Candidate models were then compared using nested model comparisons with sequential F-tests based on analysis of variance (ANOVA) to make the final model selection.

## Results

The average number of streamlines for each tract and hemisphere is reported in Table 2. When comparing the number of streamlines across hemispheres, no hemispheric lateralization was observed (β = −0.019, SE = 0.30, *t*_(237)_ = −0.061, *p* = .951).

**Table 2.** Mean (± SD) number of streamlines for each thalamo-amygdala pathway by hemisphere.

| Tract | Left hemisphere | Right hemisphere |
| --- | --- | --- |
| MGB - BLA | 418.88 (919.41) | 776.00 (2131.60) |
| PuM - BLA | 133.71 (300.17) | 220.76 (863.64) |
| PuInf - BLA | 2900.44 (3507.52) | 2185.44 (3905.60) |
| PuL - BLA | 9.21 (15.22) | 64.59 (156.82) |

Abbreviations: MGB = medial geniculate body, BLA = basolateral amygdala, PuM = medial pulvinar, PuInf = inferior pulvinar, PuL = lateral pulvinar. No hemispheric lateralization of streamline counts was observed across pathways ( $\beta = -0.019$ , $SE = 0.30$ , $t_{(237)} = -0.061$ , $p = .951$ )

### No relationship between thalamo-amygdala fiber density and trait anxiety

When examining the relationship between fiber density of the thalamo-amygdala tracts and trait anxiety scores, we fitted a linear regression model including all eight thalamo-amygdala pathways as predictors of trait anxiety (STAI-T). This full model did not reach statistical significance. PCA revealed substantial shared variance across fiber-density measures of the tracts, indicating redundancy among predictors. Nested model comparisons indicated that adding additional predictors did not significantly improve model fit relative to simpler models (all p-values > .20; see Table 3). Although the model including only the right PuInf tract yielded the lowest model complexity, it did not reach statistical significance, and no thalamo-amygdala pathway showed a reliable association with trait anxiety.

**Table 3.** Sequential ANOVA model comparison for trait anxiety (STAI-T)

| Model | Added Predictors | Res. *df* | RSS | $\Delta df$ | $\Delta SS$ | *F* | *p* |
| --- | --- | --- | --- | --- | --- | --- | --- |
| 1 | Right PulInf | 32 | 18529 | - | - | - | - |
| 2 | Model 1 + Right PuM | 31 | 17286 | 1 | 1243.23 | 1.87 | .183 |
| 3 | Model 2 + Left MGB | 30 | 16989 | 1 | 296.21 | 0.45 | .510 |
| 4 | Model 3 + Right MGB | 29 | 16825 | 1 | 164.10 | 0.25 | .623 |
| 5 | Model 4 + Left PulInf | 28 | 16665 | 1 | 160.71 | 0.24 | .627 |
| 6 | Model 5 + Left PuL | 27 | 16606 | 1 | 58.29 | 0.09 | .769 |
| 7 | Model 6 + Left PuM | 26 | 16597 | 1 | 9.48 | 0.01 | .906 |
| 8 | Full model | 25 | 16592 | 1 | 5.12 | < 0.01 | .931 |

Note: Models were compared sequentially using Type I sums of squares. Abbreviations: Res. *df* = Residual degrees of freedom, RSS = Residual Sum of Squares, $\Delta df$ = change in degrees of freedom, $\Delta SS$ = change in residual sum of squares relative to the previous model, *F* = *F*-statistic, *p* = *p*-value, MGB = medial geniculate body, BLA = basolateral amygdala, PuM = medial pulvinar, PulInf = inferior pulvinar, PuL = lateral pulvinar. Adding additional predictors did not significantly improve model fit at any step.

### Fiber density of left thalamo-amygdala tracts was linearly associated with state anxiety

To examine the relationship between fiber density of the thalamo-amygdala tracts and state anxiety scores, we initially fitted a linear regression model including all eight thalamo-amygdala pathways as predictors of state anxiety (STAI-S). As in the case of trait anxiety, this full model did not reach statistical significance. PCA again revealed substantial covariance across tract fiber-density measures, indicating redundancy among predictors. Nested model comparisons revealed that the model including three left thalamo-amygdala pathways (i.e., left MGB, left PuM and left PuInf) was the only candidate model that significantly improved model fit relative to simpler models (*F*(1,30) = 4.92, *p* = .036), whereas inclusion of additional pathways did not provide further explanatory benefit (see Table 4). Based on these model comparisons, the three-pathway model was retained as the most parsimonious explanatory model for subsequent analyses.

**Table 4.** Sequential ANOVA model comparison for state anxiety (STAI-S)

| Model | Added Predictors | Res. *df* | RSS | $\Delta df$ | $\Delta SS$ | *F* | *p* |
| --- | --- | --- | --- | --- | --- | --- | --- |
| 1 | Left MGB | 32 | 12261.3 | - | - | - | - |
| 2 | Model 1 + Left PuInf | 31 | 11121.7 | 1 | 1139.52 | 3.44 | .076 |
| 3 | Model 2 + Left PuM | 30 | 9492.9 | 1 | 1628.89 | 4.92 | **.036** |
| 4 | Model 3 + Right PuL | 29 | 8805.2 | 1 | 687.63 | 2.08 | .162 |
| 5 | Model 4 + Right PuM | 28 | 8678.4 | 1 | 126.79 | 0.38 | .542 |
| 6 | Model 5 + Right PuInf | 27 | 8522.8 | 1 | 155.64 | 0.47 | .499 |
| 7 | Model 6 + Right MGB | 26 | 8373.0 | 1 | 149.82 | 0.45 | .507 |
| 8 | Full model | 25 | 8279.7 | 1 | 93.31 | 0.28 | .600 |

Note: Models were compared sequentially using Type I sums of squares. Abbreviations: Res. *df* = Residual degrees of freedom, RSS = Residual Sum of Squares, $\Delta df$ = change in degrees of freedom, $\Delta SS$ = change in residual sum of squares relative to the previous model, *F* = *F*-statistic, *p* = *p*-value, MGB = medial geniculate body, BLA = basolateral amygdala, PuM = medial pulvinar, PuInf = inferior pulvinar, PuL = lateral pulvinar. Only model 3 significantly improved model fit (*p* < .05).

After confirming all the model assumptions were met (i.e., normality of residuals, absence of problematic multicollinearity, homoscedasticity and no autocorrelation), the selected model significantly explained variance in state anxiety, *F*(3,30) = 4.50, *p* = .010, accounting for 24.1% of the variance (adjusted R^2^ = .241; R^2^ = .310). Within this model, higher STAI-S scores were associated with lower fiber density in left MGB-BLA (β = −8.69, *t* = −2.63, *p* = .013) and left PuM-BLA (β = −8.57, *t* = −2.27, *p* = .031), whereas higher fiber density in left PuInf-BLA was associated with higher state anxiety (β = 12.84, *t* = 2.81, *p* = .009). The tract-specific associations and corresponding 3D tract reconstructions are shown in Fig. 2.

**Figure 2.**
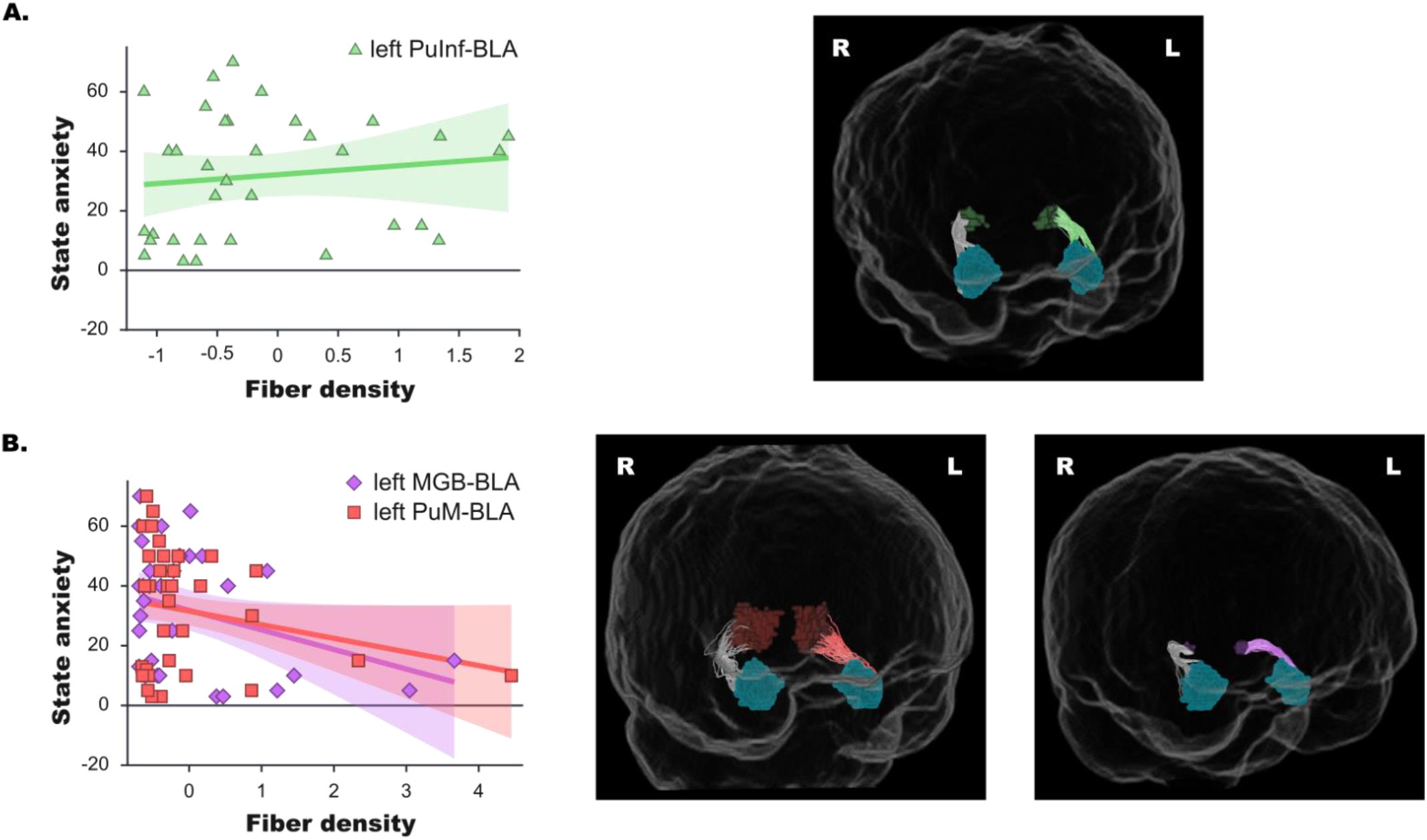
Tract-specific associations between thalamo-amygdala fiber density and state anxiety. Abbreviations: L = left, R = right, PuInf = inferior pulvinar, BLA = basolateral amygdala, MGB = medial geniculate body, PuM = medial pulvinar. (A left) Positive association between fiber density in the left inferior pulvinar-basolateral amygdala (PuInf-BLA) tract and state anxiety scores (STAI-S). Each triangle represents an individual participant. The solid line indicates the fitted linear regression, with the shaded area denoting the 95% confidence interval. (A right) 3D probabilistic tractography reconstruction of the corresponding tract in a representative subject. Fibers are seeded from the PuInf (dark green) and terminate in the BLA (petrol), with streamlines shown in light green. (B left) Negative associations between fiber density in the left medial geniculate body-basolateral amygdala (MGB-BLA) and left medial pulvinar-basolateral amygdala (PuM-BLA) tracts and state anxiety scores. Each symbol (diamond for MGB-BLA and square for PuM-BLA) represents an individual participant, with separate regression lines and confidence intervals for each tract. (B right) 3D probabilistic tractography reconstruction of the corresponding tracts in a representative subject. Fibers are seeded from the MGB (dark purple) and PuM (light red) and terminate in the BLA (petrol), with streamlines shown in light purple and light red respectively. Streamlines shown in grey represent right-hemisphere fibers, which did not contribute to the final model. Scatterplots created with BioRender; 3D probabilistic tractography reconstructions were generated using MRtrix.

To further assess the hemispheric specificity of these effects, we conducted additional control analyses by fitting linear regression models including only right-hemispheric pathways, either as a full set (four tracts) or as the corresponding subset of three pathways (right MGB, right PuM and right PuInf). None of these models significantly explained variance in state anxiety (all p-values > .554), indicating that the observed associations were specific to left thalamo-amygdala pathways.

Together, these findings suggest differentiated associations between left thalamo-amygdala pathways and state anxiety, with left MGB-BLA and PuM-BLA tracts showing a convergent negative relationship with anxiety, and the left PuInf-BLA pathway exhibiting the opposite pattern.

## Discussion

Our results demonstrate that the structural integrity of three left-hemispheric thalamo-amygdala pathways was associated with individual differences in state anxiety. Specifically, the fiber density of pathways connecting the basolateral amygdala (BLA) with the left medial geniculate body (lMGB-BLA), left medial pulvinar (lPuM-BLA) and left inferior pulvinar (lPuInf-BLA) collectively explained 24.1% of the variance in state anxiety scores. These findings indicate a robust association between left thalamo-amygdala pathway structure and inter-individual differences in state anxiety, suggesting possible hemispheric asymmetries in the functional specialization of the “low roads”.

### State, but not trait, anxiety was predicted by thalamo-amygdala fiber density

In the present study, although trait and state anxiety dimensions were positively correlated with each other (ρ = .46, *p* = .006), individual fiber density of three left thalamo-amygdala pathways was selectively associated with state anxiety, whereas no such association was observed for trait anxiety in either hemisphere.

Trait anxiety represents a stable, long-term personality dimension, whereas state anxiety reflects a more transient affective response to an immediate threat [Saviola et al., 2020]. The thalamo-amygdala pathways investigated here, previously linked to fear-related processing, may constitute subcortical “low roads” that support rapid threat detection by providing fast sensory input to the amygdala [Celeghin et al., 2015; Domínguez-Borràs et al., 2025; Kosteletou-Kassotaki et al., 2026a; Kragel et al., 2021; McFadyen et al., 2019]. It is therefore neurobiologically plausible that these routes are more strongly coupled with momentary anxious states, associated with transient sympathetic activity [Saviola et al., 2020] than with more stable, global personality traits. Supporting this, Bishop et al., [2007], demonstrated a clear neural dissociation, with higher state anxiety associated with increased left amygdala responses to threat, while trait anxiety was associated with reduced cortical activity. This pattern suggests that state and trait anxiety may rely on partly dissociable neural mechanisms, one more closely linked to immediate threat evaluation (state anxiety) and the other to top-down regulation (trait anxiety).

Notably, although structural changes may be more commonly associated with trait anxiety than with transient anxiety states [Saviola et al., 2020], our findings indicate that thalamo-amygdala structural strength is related to self-reported state anxiety. Given the positive correlation between state and trait anxiety in our sample, together with evidence linking amygdala structure to both anxiety dimensions [Blackmon et al., 2011], thalamo-amygdala fiber density may preferentially relate to transient affective states shaping anxiety expression. Additionally, because trait anxiety scores exhibited a broader distribution than state scores (Fig. 1), the absence of significant associations with trait anxiety may not be explained by a reduction in variability for this trait dimension. Rather, this further supports a selective relationship between thalamo-amygdala connectivity and state anxiety. Future work in clinical anxiety populations will nevertheless be essential to clarify the contribution of these pathways to both state and trait anxiety.

### Functional dissociation across thalamo-amygdala tracts: negative versus positive predictors of state anxiety

A notable aspect of our findings is the divergent relationship observed between thalamo-amygdala pathways and state anxiety. While the lMGB-BLA and lPuM-BLA pathways showed negative associations with anxiety, the lPuInf-BLA pathway showed a positive association (Fig. 2), suggesting that thalamo-amygdala connectivity may not represent a unitary affective system. Although the negative associations should be interpreted cautiously given the dispersion of individual values (Fig. 2), the overall pattern remained consistent across models and control analyses.

One possible interpretation is that reduced structural capacity or efficiency of the lMGB-BLA and lPuM-BLA pathways, associated with higher anxiety levels, may reflect less effective processing or filtering of emotionally salient signals. In contrast, the positive association observed for the lPuInf-BLA pathway may point to partially distinct contributions related to salience or threat sensitivity.

On the other hand, these differences may also relate to the distinct sensory profiles previously associated with these pathways. Whereas inferior pulvinar and its connections with BLA have been more strongly linked to affective visual processing [Froesel et al., 2021; Froesel et al., 2024; Kosteletou-Kassotaki et al., 2026a; McFadyen et al., 2019], MGB-BLA pathways appear to be preferentially associated with auditory processing [Cinca-Tomás et al., 2025; Kosteletou-Kassotaki et al., 2026a; LeDoux et al., 1990; Shinonaga et al., 1994], while medial pulvinar subregions may in turn contribute to auditory and multisensory integration [Bourgeois et al., 2020; Froesel et al., 2021; Froesel et al., 2024; Vittek et al., 2023].

In addition, previous work has reported increased pulvinar-amygdala circuitry engagement and pulvinar activity in phobic individuals [Ipser et al., 2013; Nakataki et al., 2017] although other studies suggest that enhanced pulvinar-amygdala connectivity may also be associated with reduced amygdala responses to threat, reflecting a potential regulatory mechanism [Hakamata et al., 2016]. These divergent findings potentially reflect differences across pulvinar subregions that are not typically segregated in these investigations. Previous evidence has also implicated pulvinar circuitry in attentional filtering and regulation of threat-related signals [Bishop et al., 2004; Hakamata et al., 2016], while lesion and neuroimaging studies have associated basolateral amygdala circuitry with threat monitoring, attentional regulation and response inhibition, with alterations or lesions in these circuits associated with hypervigilance and abnormal threat responding [Terburg et al., 2018]. Within this framework, the negative associations observed for lMGB-BLA and lPuM-BLA may reflect pathways involved in more adaptive processing or filtering of emotionally salient information, whereas the positive association observed for lPuInf-BLA may point to partially distinct contributions related to salience processing or enhanced sensitivity to threat-related signals. However, these interpretations remain speculative and will require direct functional investigation.

Overall, these findings support the idea that thalamo-amygdala pathways may show partially dissociable contributions to state anxiety.

### Left lateralization of thalamo-amygdala associations with state anxiety

The predominant involvement of left thalamo-amygdala pathways in anxiety-related effects aligns with previous evidence linking left-lateralized amygdala structure and function to anxiety-related processes [Bishop et al., 2004; Bishop et al., 2007; Blackmon et al., 2011; Talati et al., 2015]. In the present study, a model including three left-hemispheric thalamo-amygdala pathways best explained variability in state anxiety, suggesting that left-hemispheric thalamo-amygdala circuitry may be more strongly related to transient anxiety states than equivalent right-hemispheric pathways.

This interpretation is consistent with previous findings showing altered left amygdala responses and reduced attentional influences on amygdala activity in highly anxious individuals [Bishop et al., 2004; Bishop et al., 2007], as well as structural associations of left amygdala and pulvinar volume with anxiety-related measures and symptom improvement [Blackmon et al., 2011; Talati et al., 2015]. Importantly, additional control analyses including only right-hemispheric pathways did not significantly explain variance in state anxiety (see Results). However, given the heterogeneous literature on hemispheric asymmetries on anxiety and emotional processing, future studies using larger samples and multimodal approaches will be necessary to clarify the robustness, specificity and functional significance of these lateralized associations.

## Conclusion

The present study identifies four thalamo-amygdala “low roads” in humans, previously associated with affective processing. Among them, we provide evidence that structural connectivity across three left-hemispheric thalamo-amygdala pathways is selectively associated with state, but not trait, anxiety, highlighting their potential contribution to transient affective states. Importantly, our results reveal a dissociation within thalamo-amygdala circuitry: while the left MGB-BLA and medial pulvinar-BLA pathways were negatively related to state anxiety, the left inferior pulvinar-BLA pathway showed a positive association. These effects were specific to the left-hemisphere, supporting a lateralized organization of these subcortical pathways. Together, these findings suggest that thalamo-amygdala connectivity does not constitute a unitary system, but rather a heterogeneous network with partially differentiated roles in affective processing, and suggest that distinct left-hemispheric pathways may contribute differently to variability in state anxiety. By highlighting both pathway-specific and hemispheric differences, this work advances current models of subcortical affective function and identifies specific pathways that may contribute to individual variability in transient anxiety states. However, given the correlational and structural nature of these findings, direct functional investigations will be necessary to clarify the mechanisms underlying these associations.

## Author Contributions

**Martina T. Cinca-Tomás:** conceptualization, investigation, formal analysis, methodology, software, visualization, and writing – original draft. **Emmanouela Kosteletou-Kassotaki:** conceptualization, investigation, formal analysis, software, and writing – review and editing. **Judith Domínguez-Borràs:** conceptualization, methodology, supervision, and writing – review and editing.

## Data-availability Statement

Raw data have been deposited at OpenNeuro and is publicly available as of the date of publication. DOI: 10.18112/openneuro.ds007690.v1.0.0

## Ethics Statement

All procedures were approved by the ethics committee of the University of Barcelona (RB00003099 - CER042405), in accordance with the Declaration of Helsinki. All participants received monetary compensation for their participation.

## Acknowledgments

This work was funded by the European Union. Views and opinions expressed are, however, those of the authors only and do not necessarily reflect those of the European Union or the European Research Council Executive Agency. Neither the European Union nor the granting authority can be held responsible for them. Supported by ERC grant (HumanSUBthreat, 101088954), the Spanish Ministry of Science and Innovation (PID2020-116311GA-I00), the María de Maeztu Center of Excellence (MDM-2017-07-29-20-2) and the Generalitat de Catalunya (SGR2021-00356).

## References

Bishop SJ, Duncan J, Lawrence AD (2004): State anxiety modulation of the amygdala response to unattended threat-related stimuli. J Neurosci 24:10364–10368.

Bishop SJ, Jenkins R, Lawrence AD (2007): Neural Processing of Fearful Faces: Effects of Anxiety are Gated by Perceptual Capacity Limitations. Cereb Cortex 17:1595–1603. https://academic.oup.com/cercor/article-lookup/doi/10.1093/cercor/bhl070.

Blackmon K, Barr WB, Carlson C, Devinsky O, DuBois J, Pogash D, Quinn BT, Kuzniecky R, Halgren E, Thesen T (2011): Structural evidence for involvement of a left amygdala-orbitofrontal network in subclinical anxiety. Psychiatry Res - Neuroimaging 194:296–303.

Bourgeois A, Guedj C, Carrera E, Vuilleumier P (2020): Pulvino-cortical interaction: An integrative role in the control of attention. Neurosci Biobehav Rev 111:104–113. 10.1016/j.neubiorev.2020.01.005.

Carr JA (2015): I’ll take the low road: the evolutionary underpinnings of visually triggered fear. Front Neurosci 9:414. http://www.ncbi.nlm.nih.gov/pubmed/28119645.

Celeghin A, de Gelder B, Tamietto M (2015): From affective blindsight to emotional consciousness. Conscious Cogn 36:414–425. 10.1016/j.concog.2015.05.007.

Cinca-Tomás MT, Kosteletou-Kassotaki E, Costa-Faidella J, Escera C, Domínguez-Borràs J (2025): An auditory “low road” for threat in humans sensitive to fast temporal cues. http://biorxiv.org/lookup/doi/10.64898/2025.12.24.690990.

Cordero-Grande L, Christiaens D, Hutter J, Price AN, Hajnal J V. (2019): Complex diffusion-weighted image estimation via matrix recovery under general noise models. Neuroimage 200:391–404. 10.1016/j.neuroimage.2019.06.039.

Domínguez-Borràs J, Bourgeois A, Vuilleumier P (2025): Affective Biases in Attention, Cognitive Control, and Awareness. In:. The Cambridge Handbook of Human Affective Neuroscience. Cambridge University Press. pp 385–407. https://www.cambridge.org/core/product/identifier/9781009342919%23bp25/type/book_part.

Domínguez-Borràs J, Vuilleumier P (2022): Amygdala function in emotion, cognition, and behavior. Handb Clin Neurol 187:359–380.

Etkin A, Klemenhagen KC, Dudman JT, Rogan MT, Hen R, Kandel ER, Hirsch J (2004): Individual differences in trait anxiety predict the response of the basolateral amygdala to unconsciously processed fearful faces. Neuron 44:1043–1055.

Froesel M, Cappe C, Ben Hamed S (2021): A multisensory perspective onto primate pulvinar functions. Neurosci Biobehav Rev 125:231–243.

Froesel M, Gacoin M, Clavagnier S, Hauser M, Goudard Q, Ben Hamed S (2024): Macaque claustrum, pulvinar and putative dorsolateral amygdala support the cross-modal association of social audio-visual stimuli based on meaning. Eur J Neurosci 59:3203–3223.

Hakamata Y, Sato E, Komi S, Moriguchi Y, Izawa S, Murayama N, Hanakawa T, Inoue Y, Tagaya H (2016): The functional activity and effective connectivity of pulvinar are modulated by individual differences in threat-related attentional bias. Sci Rep 6:1–12.

Iglesias JE, Insausti R, Lerma-Usabiaga G, Bocchetta M, Van Leemput K, Greve DN, van der Kouwe A, Fischl B, Caballero-Gaudes C, Paz-Alonso PM (2018): A probabilistic atlas of the human thalamic nuclei combining ex vivo MRI and histology. Neuroimage 183:314–326.

Ipser JC, Singh L, Stein DJ (2013): Meta-analysis of functional brain imaging in specific phobia. Psychiatry Clin Neurosci 67:311–322.

Janak PH, Tye KM (2015): From circuits to behaviour in the amygdala. Nature 517:284–292.

Jeurissen B, Tournier JD, Dhollander T, Connelly A, Sijbers J (2014): Multi-tissue constrained spherical deconvolution for improved analysis of multi-shell diffusion MRI data. Neuroimage 103:411–426. 10.1016/j.neuroimage.2014.07.061.

Keifer OP, Gutman DA, Hecht EE, Keilholz SD, Ressler KJ (2015): A comparative analysis of mouse and human medial geniculate nucleus connectivity: A DTI and anterograde tracing study. Neuroimage 105:53–66. 10.1016/j.neuroimage.2014.10.047.

Kellner E, Dhital B, Kiselev VG, Reisert M (2016): Gibbs-ringing artifact removal based on local subvoxel-shifts. Magn Reson Med 76:1574–1581.

Khalil V, Faress I, Mermet-Joret N, Kerwin P, Yonehara K, Nabavi S (2023): Subcortico-amygdala pathway processes innate and learned threats. Elife 12:1–26.

Koller K, Rafal RD, Platt A, Mitchell ND (2019): Orienting toward threat: Contributions of a subcortical pathway transmitting retinal afferents to the amygdala via the superior colliculus and pulvinar. Neuropsychologia 128:78–86.

Kosteletou-Kassotaki E, Cinca-Tomás MT, Varriano F, Soria G, Prats-Galino A, Domínguez-Borràs J (2026a): A Direct Auditory Subcortical Route to the Amygdala Associated with Fear in Humans. J Neurosci 46:e1431252026. https://www.jneurosci.org/lookup/doi/10.1523/JNEUROSCI.1431-25.2026.

Kosteletou-Kassotaki E, Mengxing L, Cinca-Tomás MT, Paz-Alonso PM, Domínguez-Borràs J (2026b): Mapping the subcortical pathways associated with fear in the human brain: multiple thalamo-amygdala connections revealed by high-resolution tractography. Neuroimage 335:121983. http://www.ncbi.nlm.nih.gov/pubmed/42107615.

Kragel PA, Ćeko M, Theriault J, Chen D, Satpute AB, Wald LW, Lindquist MA, Feldman Barrett L, Wager TD (2021): A human colliculus-pulvinar-amygdala pathway encodes negative emotion. Neuron 109:2404–2412.e5.

LeDoux JE, Sakaguchi A, Reis DJ (1984): Subcortical efferent projections of the medial geniculate nucleus mediate emotional responses conditioned to acoustic stimuli. J Neurosci 4:683–98. http://www.jneurosci.org/cgi/content/abstract/4/3/683%5Cnpapers://bb18545c-4283-413e-92cc-6a26fd17ba65/Paper/p69.

LeDoux JE (2000): Emotion circuits in the brain. Annu Rev Neurosci 23:155–184.

LeDoux JE, Farb C, Ruggiero DA (1990): Topographic organization of neurons in the acoustic thalamus that project to the amygdala. J Neurosci 10:1043–1054.

McFadyen J (2019): Investigating the Subcortical Route to the Amygdala Across Species and in Disordered Fear Responses. J Exp Neurosci 13:1179069519846445. http://www.ncbi.nlm.nih.gov/pubmed/31068755.

McFadyen J, Dolan RJ, Garrido MI (2020): The influence of subcortical shortcuts on disordered sensory and cognitive processing. Nat Rev Neurosci 21:264–276. 10.1038/s41583-020-0287-1.

McFadyen J, Mattingley JB, Garrido MI (2019): An afferent white matter pathway from the pulvinar to the amygdala facilitates fear recognition. Elife 8:1–51.

Nakataki M, Soravia LM, Schwab S, Horn H, Dierks T, Strik W, Wiest R, Heinrichs M, De Quervain DJF, Federspiel A, Morishima Y (2017): Glucocorticoid Administration Improves Aberrant Fear-Processing Networks in Spider Phobia. Neuropsychopharmacology 42:485–494.

Neumeister P, Feldker K, Heitmann CY, Buff C, Brinkmann L, Bruchmann M, Straube T (2018): Specific amygdala response to masked fearful faces in post-traumatic stress relative to other anxiety disorders. Psychol Med 48:1209–1217.

Pessoa L, Adolphs R (2010): Emotion processing and the amygdala: from a “low road” to “many roads” of evaluating biological significance. Nat Rev Neurosci 11:773–83. https://www.nature.com/articles/nrn2920.

Phelps EA, LeDoux JE (2005): Contributions of the amygdala to emotion processing: From animal models to human behavior. Neuron 48:175–187.

Rafal RD, Koller K (2025): Collothalamic projections to the human amygdala: hemispheric asymmetry modulates trait anxiety. J Neurophysiol 133:1054–1066.

Romanski LM, LeDoux JE (1992): Equipotentiality of thalamo-amygdala and thalamo-cortico-amygdala circuits in auditory fear conditioning. J Neurosci 12:4501–4509.

Saviola F, Pappaianni E, Monti A, Grecucci A, Jovicich J, De Pisapia N (2020): Trait and state anxiety are mapped differently in the human brain. Sci Rep 10:1–11. 10.1038/s41598-020-68008-z.

Saygin ZM, Kliemann D, Iglesias JE, van der Kouwe AJW, Boyd E, Reuter M, Stevens A, Van Leemput K, McKee A, Frosch MP, Fischl B, Augustinack JC (2017): High-resolution magnetic resonance imaging reveals nuclei of the human amygdala: manual segmentation to automatic atlas. Neuroimage 155:370–382. 10.1016/j.neuroimage.2017.04.046.

Shinonaga Y, Takada M, Mizuno N (1994): Direct projections from the non-laminated divisions of the medial geniculate nucleus to the temporal polar cortex and amygdala in the cat. J Comp Neurol 340:405–426.

Smith RE, Tournier J-D, Calamante F, Connelly A (2012): Anatomically-constrained tractography: improved diffusion MRI streamlines tractography through effective use of anatomical information. Neuroimage 62:1924–38. 10.1016/j.neuroimage.2012.06.005.

Smith RE, Tournier J-D, Calamante F, Connelly A (2015): SIFT2: Enabling dense quantitative assessment of brain white matter connectivity using streamlines tractography. Neuroimage 119:338–51. 10.1016/j.neuroimage.2015.06.092.

Smith SM, Jenkinson M, Woolrich MW, Beckmann CF, Behrens TEJ, Johansen-Berg H, Bannister PR, De Luca M, Drobnjak I, Flitney DE, Niazy RK, Saunders J, Vickers J, Zhang Y, De Stefano N, Brady JM, Matthews PM (2004): Advances in functional and structural MR image analysis and implementation as FSL. Neuroimage 23:S208–S219. http://www.ncbi.nlm.nih.gov/pubmed/15501092.

Spielberger C, Gorsuch R, Lushene R. (1982): STAI. Cuestionario de Ansiedad Estado/Rasgo. Manual 5 ed Madrid. https://books.google.es/books/about/STAI_Cuestionario_de_Ansiedad_Estado_Ras.html?id=T6CUtwAACAAJ&source=kp_book_description&redir_esc=y%0Ahttp://server1.docfoc.com/uploads/Z2016/01/21/gqkneIRwOZ/8c6c0ccbdd3ab122da807f49625f4fe2.pdf.

Steuwe C, Daniels JK, Frewen PA, Densmore M, Theberge J, Lanius RA (2015): Effect of direct eye contact in women with PTSD related to interpersonal trauma: Psychophysiological interaction analysis of connectivity of an innate alarm system. Psychiatry Res - Neuroimaging 232:162–167. 10.1016/j.pscychresns.2015.02.010.

Talati A, Pantazatos SP, Hirsch J, Schneier F (2015): A pilot study of gray matter volume changes associated with paroxetine treatment and response in social anxiety disorder. Psychiatry Res - Neuroimaging 231:279–285. 10.1016/j.pscychresns.2015.01.008.

Tamietto M, De Gelder B (2010): Neural bases of the non-conscious perception of emotional signals. Nat Rev Neurosci 11:697–709.

Terburg D, Scheggia D, Triana del Rio R, Klumpers F, Ciobanu AC, Morgan B, Montoya ER, Bos PA, Giobellina G, van den Burg EH, de Gelder B, Stein DJ, Stoop R, van Honk J (2018): The Basolateral Amygdala Is Essential for Rapid Escape: A Human and Rodent Study. Cell 175:723–735.e16. 10.1016/j.cell.2018.09.028.

Veraart J, Novikov DS, Christiaens D, Ades-aron B, Sijbers J, Fieremans E (2016): Denoising of diffusion MRI using random matrix theory. Neuroimage 142:394–406. 10.1016/j.neuroimage.2016.08.016.

Vittek A-L, Juan C, Nowak LG, Girard P, Cappe C (2023): Multisensory integration in neurons of the medial pulvinar of macaque monkey. Cereb Cortex 33:4202–4215. http://www.ncbi.nlm.nih.gov/pubmed/35662511.

Vuilleumier P (2005): How brains beware: Neural mechanisms of emotional attention. Trends Cogn Sci 9:585–594.

